# Coyotes in New York city carry variable dog genomic ancestry and influence their interactions with humans

**DOI:** 10.1101/2022.08.15.504024

**Authors:** Anthony Caragiulo, Stephen J. Gaughran, Neil Duncan, Christopher Nagy, Mark Weckel, Bridgett M. vonHoldt

## Abstract

Coyotes are ubiquitous on the North American landscape as a result of their recent expansion across the continent. They have been documented in the heart of some of the most urbanized cities, such as Chicago, Los Angeles, and New York City. Here, we explored the genomic composition of coyotes in the New York metropolitan area to investigate if genomic demography and admixture differs from expected for urban-dwelling canids. We identified moderate-to-high estimates of relatedness among coyotes living in Queens and adjacent neighborhoods, suggestive of a relatively small population. Although we found low background levels of domestic dog ancestry across most coyotes in our sample, we identified a male suspected to be a first-generation coyote-dog hybrid, as well as his two putative backcrossed offspring that carried approximately 25% dog ancestry. The male coyote-dog hybrid and one backcrossed offspring each carried two mutations that are known to increase human-directed hypersociability in dogs and gray wolves. An additional, unrelated coyote with little dog ancestry also carries two of these mutations. These genetic patterns suggest that gene flow from domestic dogs may become an increasingly important consideration as coyotes continue to inhabit metropolitan regions.

## Introduction

Coyotes (*Canis latrans*) are synanthropes (Gehrt et al. 2011) who have undergone a dramatic range expansion across much of North and Central America. Prior to 1900, coyotes were restricted to the western two-thirds of North America (Hody & Kays 2018; Hinton et al 2019). They subsequently expanded into eastern North America likely due to various interacting factors, that include the extirpation of apex predators (e.g. wolves, cougars) and the conversion of once-forested landscapes to agricultural landscapes, which provided ample opportunity for coyotes to hybridize with eastern wolves (*C. lycaon*), red wolves (*C. rufus*), gray wolves (*C. lupus*), and domestic dogs (*C. familiaris*) (Lehman et al. 1991; Way et al. 2010; Kays et al. 2009; Kays et al. 2010; vonHoldt et al. 2011; Benson et al. 2012; Wheeldon et al. 2013; Monzon et al 2014; Bohling and Waits 2015; Gese et al. 2015; vonHoldt et al. 2016; Hody and Kays 2018; vonHoldt and Aardema 2020). Gene flow between these canids is a possible mechanism by which new, adaptive genetic variation may have promoted colonization and survival in eastern habitats (Kays et al 2010, vonHoldt et al 2011; Thornton and Murray 2014; Hody & Kays 2018; Heppenheimer et al 2018c). Further, coyotes have been documented in nearly every habitat type and landscape in North America, encompassing both natural and disturbed habitats (Gehrt et al 2009; Weckel et al 2015), including major cities such as Los Angeles, Chicago, and most recently, New York City (NYC) (Gompper 2002; Gehrt et al 2009; Nagy et al. 2016; Hody and Kays 2018; Henger et al. 2020; Bradfield et al. 2022).

NYC is the most densely populated city in the United States with over 8.8 million people and approximately 27,000 people per square mile (United States Census Bureau 2020). The first documented evidence of a coyote in the NYC metropolitan area occurred in 1994 in the Bronx (Toomey et al. 2012). Another coyote was captured in Central Park in 1999 (Martin 1999). The NYC metropolitan area coyote population has since increased and become well established (Henger et al. 2020). A previous genetic survey of the NYC metropolitan area revealed that the NYC coyote population is genetically differentiated from populations in adjacent states, and with other aspects of genetic parameters, the NYC coyote inhabitants likely descended from a small founder population with limited flow of new genetic variation into the urban ecosystem (DeCandia et al. 2019, Henger et al. 2020).

Urban areas present a unique ecosystem with many ecological opportunities for coyotes. Among the *Canis* species, coyotes have a history of being characterized as generalists and opportunistic predators with highly variable diets (Bekoff 1977; MacCracken and Hansen 1987; Windberg and Mitchell 1990; Arjo et al. 2002). Natural prey species can be retained in remnant yet fragmented heterogeneous urban patches, such as parks, and their diet is likely to be supplemented with anthropogenic food subsidies (Nagy et al 2016; Henger et al 2019; Duncan et al 2020; Larson et al 2020, Bradfield et al. 2022; Henger et al. *in press*). Coyotes are an ideal system in which to study range expansion and urban ecological dynamics since they have the combined influence of their generalist ecology and a demographic history of introgression. Historic and ancient hybridization among North American canids has been well described (reviewed in vonHoldt and Aardema 2020). Despite this extensive body of research, there has infrequent genetic detection of first (F1) or second (F2) generation coyote-dog hybrids in urban environments, or their recent backcrosses, even when physical and behavioral phenotypes suggest possible introgression (Mowry et al. 2021). Here, we use a reduced representation method to collect genome-wide genotype data to probe the genomic ancestry of coyotes in NYC and the surrounding areas, including coyotes suspected to represent a family group with coyote-dog hybrids living in a small greenspace in the Ditmars/East Elmhurst area of Queens, NYC, near Elmjack Baseball Field (40.7765° N, 073.8901° W WGS84 - hereafter referred to as Elmjack). We assessed their genetic relatedness and compared genetic patterns to coyotes sampled in an adjacent region on Long Island (New York). We inferred local genetic ancestry for each individual and explored how variation in functional genes may contribute to the behavioral phenotype of the Elmjack family.

## Methods

### Sample collection and genomic DNA preparation

We obtained 16 blood or tissue samples from coyote individuals in NYC and surrounding areas (Table S1), including a suspected pack near Elmjack Baseball Field in Queens (Fig. 1). This work was conducted under the approved Princeton University IACUC protocol 1961A. We obtained high molecular weight genomic DNA using the Qiagen High Molecular Weight DNA Kit following the manufacturer’s protocol for enucleated whole blood and frozen tissue. We quantified DNA concentration using the Qubit 2.0 fluorometer system and subsequently standardized each sample’s DNA concentration to 5ng/µL.

**Figure 1.**
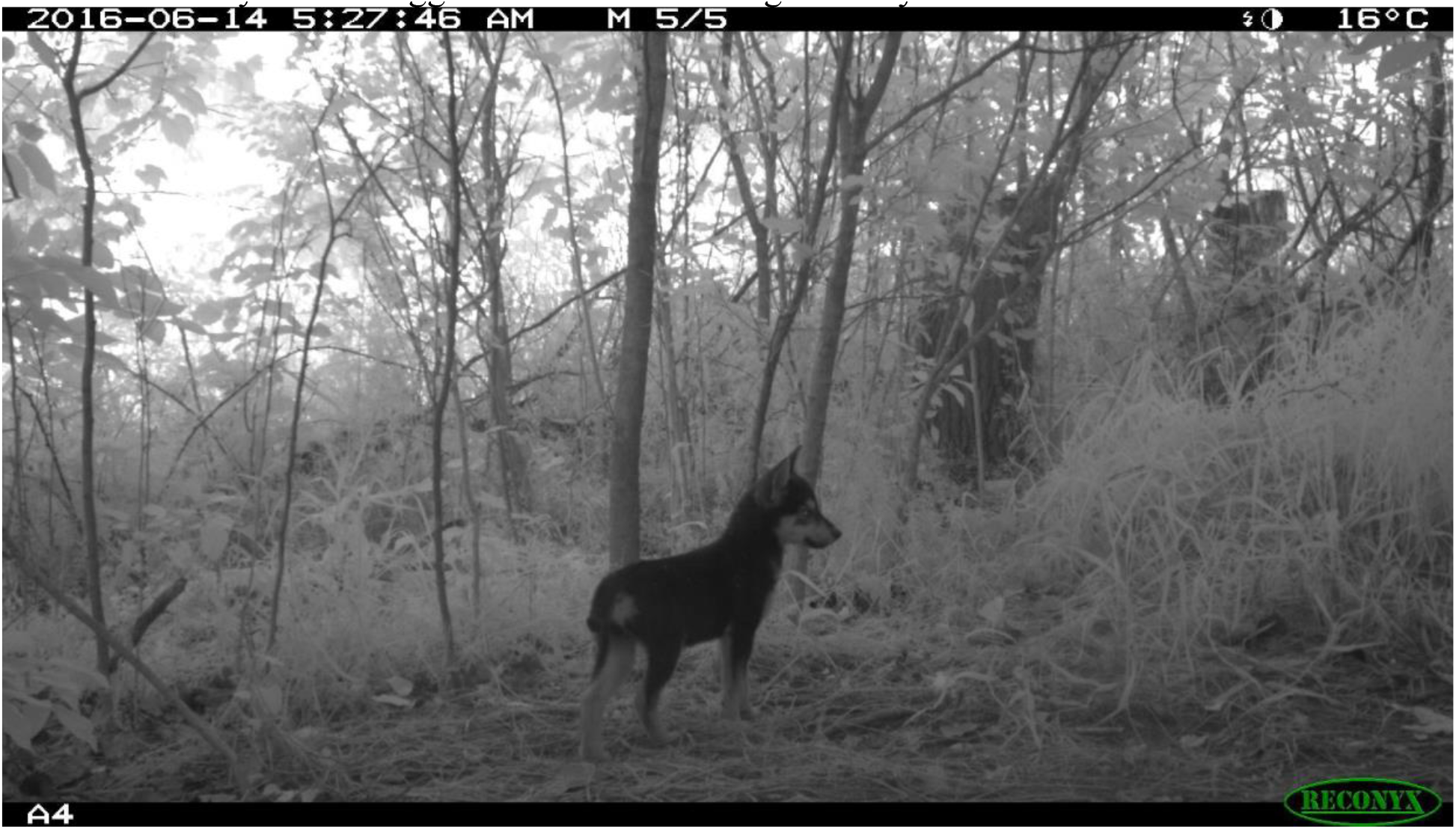
Trail camera photograph of a pup in the Elmjack coyote pack. Pelage pattern is unusual for a coyote and suggestive of domestic dog ancestry.

### RAD sequencing and bioinformatic processing

We prepared genomic DNA for restriction-site associated DNA sequencing (RADseq-capture; Ali et al. 2015) and digested genomic DNA with the *SbfI* restriction enzyme with subsequent ligation of a unique 8-bp barcoded biotinylated adapter. The barcoded adapters allowed us to then pool equal amounts of 48 DNA samples, which were then randomly sheared to 400bp in a Covaris LE220. These sheared pools were then enriched for the adapter ligated fragments using a Dynabeads M-280 streptavidin binding assay. We then prepared these enriched pools for Illumina NovaSeq paired-end (2×150nt) sequencing at Princeton University’s Lewis Sigler Genomics Institute core facility using the NEBnext Ultra II DNA Library Prep Kit. We used Agencourt AMPure XP magnetic beads for any library purification step and the size selection for fragments 300-400bp in size.

We processed the sequence data by retaining the read (and its pair) that contained the unique barcode and the remnant *SbfI* recognition motif using a custom perl script. We then further processed these reads in *STACKS* v2 (Catchen et al. 2013; Rochette et al. 2019). We first rescued specific barcoded reads using the *process_radtags* module with a 2bp mismatch and retained reads with a quality score >9. We used the *clone_filter* module to remove PCR duplicates for mapping to the dog genome CanFam3.1 assembly (Lindblad-Toh et al. 2005) using *bwa-mem* (Li 2013). We excluded mapped reads with MAPQ<20 and then converted the SAM files to BAM format in *Samtools* v0.1.18 (Li et al. 2009). We included 92 publicly available canid samples already mapped to the same reference genome assembly following the same methods (Table S1).

### SNP discovery and filtering

We merged the target 16 coyotes with public RADseq data from 40 reference samples representing domestic dogs, western coyotes, eastern wolves, and gray wolves prepared following the same methods (references for all public samples found in Table S1). We retained samples with a minimum of 300,000 mapped reads, which were used to construct a catalog of all polymorphic sites possible. We implemented the *gstacks* and *populations* modules in *STACKS* v2 following the recommended pipeline for data mapped to a reference genome. To increase the stringency of SNP annotation, we increased the minimum significance threshold in *gstacks* and used the marukilow model flags --vt-alpha and --gt-alpha with *p*=0.01. All SNPs discovered per locus were reported for downstream filtering. We used *VCFtools* v0.1.17 (Danecek et al. 2011) to exclude singleton and private doubleton alleles, to remove loci with more than 90% missing data across all samples, and to remove individuals with more than 20% missing data. We filtered to exclude sites with a minor allele frequency (MAF<0.03) and allowed up to 80% genotyping rate per locus (--*geno* 0.2) in *PLINK* v1.90b3i (Chang et al. 2015). For estimating pairwise relatedness coefficients in the R package *related* (Pew et al. 2015; vonHoldt et al. 2020), we constructed a “statistically neutral and unlinked” dataset of SNPs by excluding sites within 50-SNP windows that exceeded genotype correlations of *r*=0.2 (--*indep*-*pairwise* 50 5 0.2; a proxy for linkage disequilibrium or LD), significantly deviated from Hardy-Weinberg Equilibrium (HWE) with the argument --*hwe* 0.001, and increased the MAF threshold to 0.20. We used the coancestry function, the dyadic likelihood estimator (dyadml=1; Milligan 2003), and permitted inbreeding (allow.inbreeding=TRUE) to estimate relatedness coefficients.

### Sex inference

We included the Y chromosome (KP091776.1; Li et al. 2013) with the CanFam3.1 reference assembly to enable bioinformatic inference of sex. We estimated the number of reads that aligned to each Y-chromosome nucleotide for each RADseq sample. The expectation is that males will have a significantly higher number of reads aligned to the Y chromosome than females. However, there is also some variation in female-mapped reads to the Y chromosome’s pseudoautosomal region (PAR) that pairs with the X chromosome (1bp-6.7Mb; Raudsepp et al. 2012). We included 37 positive controls where field sex was reported (n males=14, females=23) (Table S1).

### Inference of autosomal canid ancestry

To infer ancestry proportions for the queried New York coyote samples, we selected a set of reference populations based on previous genome-wide studies that identified populations of little to no admixture, as well as incorporating pre-expansion demographics (Heppenheimer et al. 2018a,d; Heppenheimer et al. 2020). The four possible representative ancestral populations (n=10 each) were: western coyote, gray wolves, eastern wolves, and domestic dogs (Table S1). We included representatives of domestic dogs (*C. familiaris*) collected from North America given the possibility of recent interbreeding with dogs in an urban community. Further, we included both the eastern and gray wolves given their different known demographic histories (Heppenheimer et al. 2018d). We inferred local ancestry of the query coyotes from New York with respect to four reference populations (dog, western coyote, gray wolf, and eastern wolf) and using the SNP dataset filtered only for MAF and missingness (defined in Table S1). We implemented a two-layer hidden Markov model in the program *ELAI* (Guan 2014). The first step is for the model to evaluate LD and return a per-SNP allele dosage score that estimates the most likely ancestry and its state (heterozygous or homozygous). We opted to discard a SNP if it was missing from one of the populations. We defined the number of upper-layer clusters (-C) to be the number of references used and the lower-layer clusters (-c) to be twice the value of the -C value. Given our lack of *a priori* knowledge about potential gene flow, we analyzed four time points since admixture (-mg): 5, 10, 15, and 20 generations ago. We further implemented *ELAI* three times serially for each -*mg* parameter value with 30 EM steps. We obtained the final estimates by averaging each result from the 12 independent analyses.

### Human-directed hypersociability genotypes

The Elmjack pack established a den in a 14-acre undeveloped site in northern Queens, but after that site was developed into a parking lot for LaGuardia Airport staff, the coyotes became frequent visitors to the adjacent Elmjack Baseball Field. This field was surrounded by a small but dense strip of woods and scrubland that had grown over old construction fill and provided refugia for the coyotes during the day. This family group also used the grounds of a nearby water treatment plant and an island that housed a NYC jail (connected to the main Queens landmass by a 1.3 km bridge). Researchers received reports from residents, NYC employees, and New York State officials describing evidence that these coyotes were being fed by people both directly (i.e., direct handouts, piles of dog food, etc.) and indirectly (garbage/litter and cat food intentionally left for feral cats) at the LaGuardia Airport parking lot, the parking lot for the bridge, and the water treatment plant. Stomach content analysis after euthanasia confirmed these individuals were fed anthropogenic items (e.g., concession stand food such as hot dogs).

Because of these reports, we genotyped three loci known to be informative about human-directed hypersociability behavior in canines: Cfa6.6, Cfa6.7, and Cfa6.66 located on canine chromosome 6 (2,031,491-7,215,670bp of canfam3.1 assembly; see for details of each locus: vonHoldt et al. 2017; Tandon et al. 2019). The alleles at each locus associated with canine social behavior is a short transposable element (TE) insertion of the LINE or SINE family of elements. We used a previously developed PCR assay to obtain codominant genotypes for the TE: lacking (0 copies), heterozygous (1 copy), or homozygous (2 copies) for the TE (vonHoldt et al. 2017). PCR products were visualized and genotyped on a 2% agarose gel. Previous research has reported that higher copy number of TE insertions is significantly associated with increased social behavior directed at humans via prolonged durations of canine-human interactions (vonHoldt et al. 2017).

We explored the ancestry structure across chromosome 6, which houses the TE alleles associated with human-directed canine hypersociability. Ancestry blocks were defined by a minimum of three contiguous SNPs of the same locally inferred ancestry identity. We annotated the genes contained within each ancestry block using the UCSC Genome Browser’s Table Browser function for dog reference genome CanFam3.1 (Karolchik et al. 2004). We used the *intersect* function of *BedTools* v2.28 to annotate each ancestry block (Quinlan and Hall 2010).

## Results

### Sequencing and sex inference

We sequenced 16 target coyotes collected from the New York City metropolitan area and discovered SNPs with the inclusion of 40 additional reference genomes for ancestry analysis. We established a catalog of 2,620,950 loci with an effective per-sample sequence coverage of 8.7-fold. Four samples were excluded from all downstream ancestry inferences due to missing data (two target coyotes and two gray wolves). After filtering SNPs for MAF and missingness, we retained 53 canids (38 references and 15 target) and 26,763 SNP loci for ancestry inference. An initial check of samples was completed using PCA with 16,355 loci after excluding SNPs that significantly deviated from HWE and were highly correlated with other loci.

We aligned all 57 RADseq samples to the Y chromosome (KP091776.1; Li et al. 2013). Of the 37 animals of known sex, we report a 94.6% concordance rate, with only two mismatches. Males had a significantly higher number of reads that aligned to the Y chromosome than females (mean males=22196.2, females=1510.4, 1-tailed *t*-test of unequal variance *p*=5.18×10^−9^) (Table S1). We then inferred seven females and 13 males for the remainder of the samples that lacked observation and corrected the two mismatches. For the 17 query coyotes, we inferred this sample set to be composed of seven females and 10 males.

### Dog ancestry detected in coyotes in the New York Metro Area

The PCA revealed that the target coyotes likely contained admixed genomic contributions from both coyotes and domestic dogs (Fig. 2). We then inferred ancestry with respect to two reference groups (domestic dogs and western coyotes) and found that the target coyotes from the New York metro area carried an average of 12.5% dog ancestry (Table 1). However, this is predominantly due to three coyotes (male NY01, male NY04, and male NY05) with very high dog ancestry (46.5%, 24.5%, and 29.2% respectively). The remaining coyotes carried an average of 6.9% dog ancestry. The coyotes with high dog ancestry, and coyote T211, originated from a single location and possibly represent a family group (Fig. 3A).

**Table 1.** Autosomal ancestry proportions for each of the 15 target coyotes sampled in the New York metro area and inferred from 26,763 SNP genotypes across two reference populations (domestic dog, Cfa; western coyote, Cla).

| Sample ID | Cfa | Cla |
| --- | --- | --- |
| PC16 | 0.084 | 0.917 |
| MEW005 | 0.081 | 0.919 |
| NY04 | 0.245 | 0.755 |
| MEW001 | 0.058 | 0.942 |
| T211 | 0.094 | 0.907 |
| MEW009B | 0.048 | 0.952 |
| MEW004 | 0.092 | 0.908 |
| SH002 | 0.070 | 0.930 |
| NY01 | 0.465 | 0.535 |
| NY05 | 0.292 | 0.708 |
| MEW009T | 0.043 | 0.957 |
| MEW003 | 0.067 | 0.934 |
| SH001 | 0.125 | 0.875 |
| MEW002 | 0.056 | 0.944 |
| 51148 | 0.056 | 0.944 |

**Figure 2.**
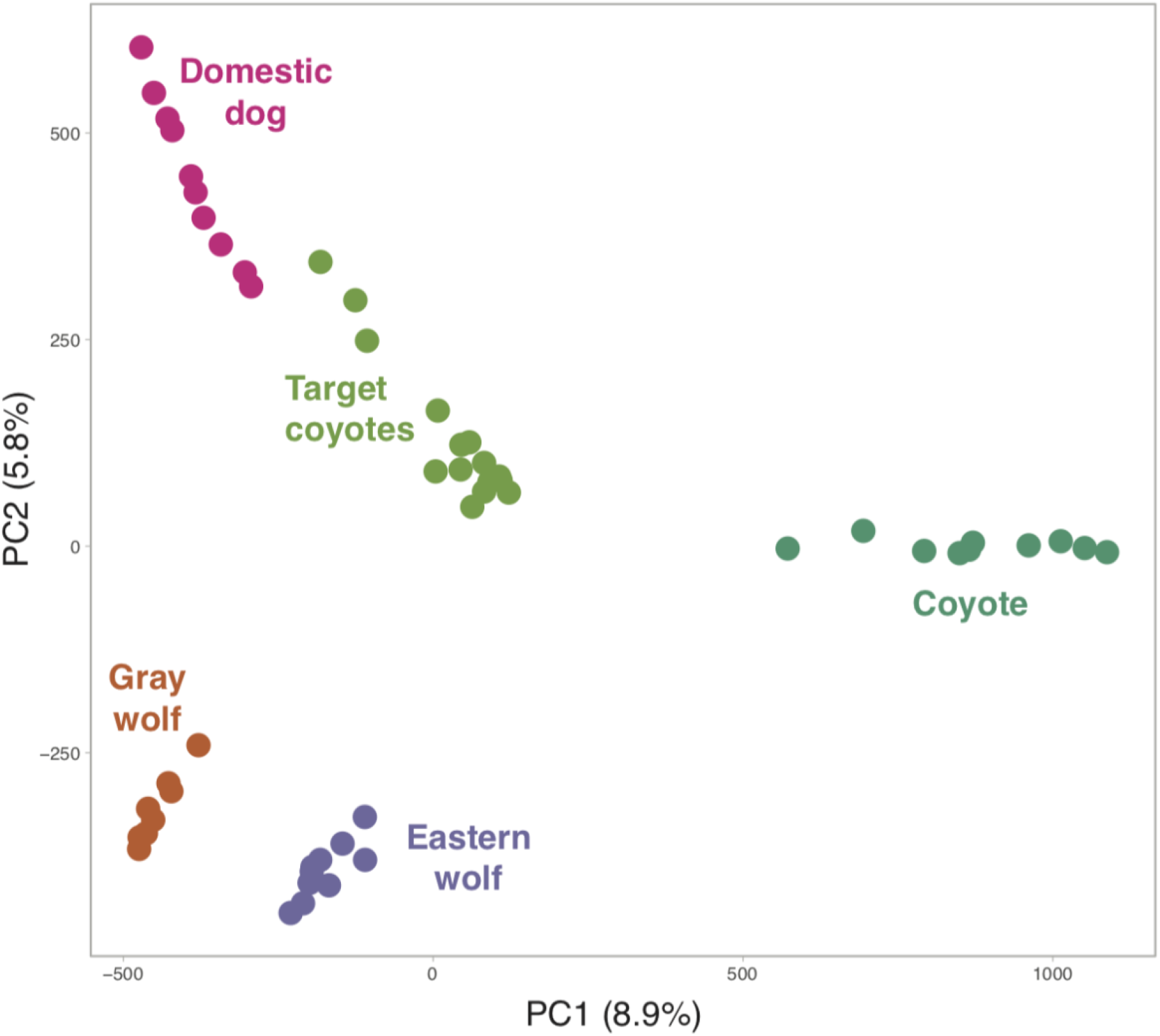
PCA of 53 canids genotyped at 16,355 unlinked and neutral SNP loci. The percent of variation explained by each axis is provided in parentheses. The 15 target coyotes were sampled in the New York City metropolitan area.

**Figure 3.**
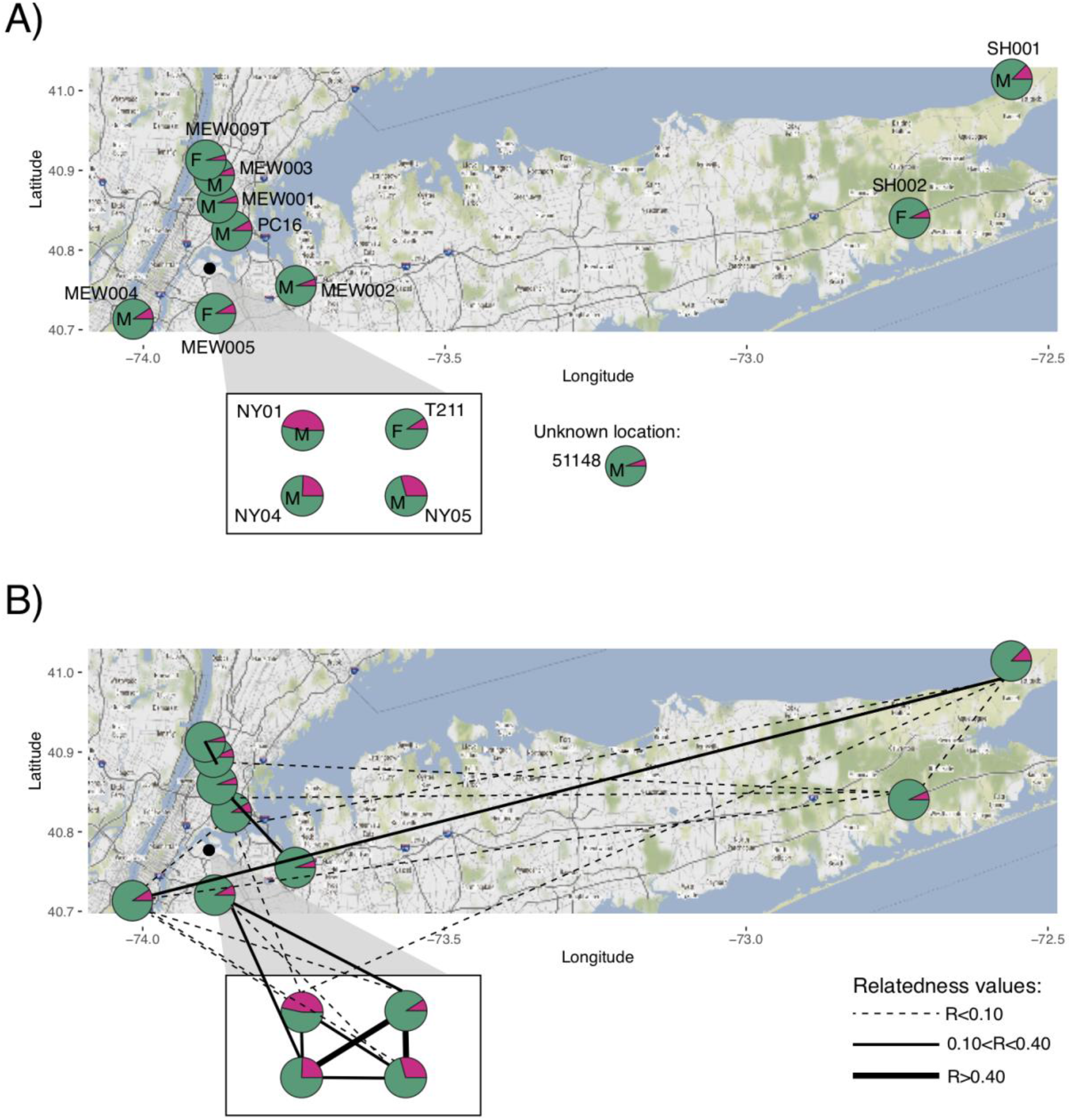
**A)** Ancestry inference for autosomal ancestry proportions inferred from 26,763 SNP loci and two reference populations (pink, domestic dog; green, western coyote) with sex indicated on each pie chart (F, female; M, male); and **B)** genetic relatedness coefficients estimated from 240 SNP loci genotyped in target coyotes sampled in the NYC metropolitan area. Inset shows four coyotes in a single location. Sample MEW009B was not plotted. Samples in panel **B)** follow the same labels found in panel **A)**.

### Family group contains F1 and F2 dog-coyote hybrids

We estimated pairwise relatedness coefficients with the dyadic likelihood estimator across 240 highly filtered SNPs genotyped in 16 target coyotes sampled in the New York metro area. We found a low average level of relatedness across 105 dyads (r=0.05) with substantial variation (s.d.=0.14) and 76 of these comparisons were unrelated (r=0) (Fig. S1; Table S2). We confirmed the duplicate sample MEW009B and MEW009T is derived from the same individual female coyote. We found the suspected family group (female T211 and three males NY01, NY04, and NY05) to have a high average relatedness (r=0.35). There was one unrelated dyad (female T211 and male NY01) with all other intra-group estimates indicative of siblings/parent-offspring or half-siblings (r=0.35-0.5) (Fig. 3B, Table 2). Spatial patterning of both ancestry and relatedness further reveals the limited dog hybridization and introgression. We found that high dog ancestry was restricted to the single pack in Elmjack, Queens, which also contained the highest inter-individual relatedness (Fig. 3B). Few other pairs had notable relatedness values (r>0.10). We found that most notable relatedness pairs were spatially adjacent (e.g. female MEW009T and male MEW003, r=0.36; female MEW005 and female T211, r=0.35). A single long-distance dyad was found between male MEW004 and male SH001, collected on opposite ends of Long Island (r=0.13) (Fig. 3B).

**Table 2.** Pairwise relatedness (r) values from the dyadic likelihood (dyadml) estimator of the putative family group of coyotes sampled in the New York metro area genotyped for 240 SNP <u>loci genotyped.</u>

| Sample ID 1 | Sample ID 2 | Dyadml $r$ |
| --- | --- | --- |
| T211 | NY05 | 0.50 |
| NY04 | T211 | 0.47 |
| NY04 | NY05 | 0.40 |
| NY01 | NY05 | 0.39 |
| NY04 | NY01 | 0.35 |
| T211 | NY01 | 0.0 |

### Human-directed hypersociability genotypes

Given the preponderance of notable dog ancestry in several of the coyotes analyzed, we followed up with a genotyping assay for TE insertions known to increase human-directed hypersocial behavior (vonHoldt et al. 2017). We found that three of the 16 coyotes genotyped carried a TE in the heterozygous genotypic state (Table 3). Of those three, two (NY01 and NY04) were previously observed to have interactions with the local human community. Based on our assessment of relatedness between NY01 (male) as a putative father of NY04 (male), we see further corroboration that both individuals are heterozygous for TE at locus Cfa6.6 and Cfa6.66 (Table 3). Coyote T211 (putative mother) and NY05 (sibling of NY04) lacked all possible TE insertions assayed.

**Table 3.** Number of transposable element insertions associated with canine human-directed hypersociability per locus. Missing data is indicated by “-”. The asterisk indicates coyotes that <u>had been documenting interacting with humans.</u>

| Sample ID | Cfa6.6 | Cfa6.7 | Cfa6.66 |
| --- | --- | --- | --- |
| MEW007 | 0 | 0 | 0 |
| PC16 | 0 | 0 | 0 |
| MEW008 | 0 | 0 | 0 |
| MEW005 | 0 | 0 | 0 |
| NY04* | 1 | 0 | 1 |
| MEW001 | 1 | 1 | 0 |
| T211* | 0 | 0 | 0 |
| MEW009B | 0 | 0 | 0 |
| MEW004 | 0 | 0 | 0 |
| SH002 | 0 | 0 | 0 |
| NY01* | 1 | 0 | 1 |
| NY05* | 0 | 0 | 0 |
| MEW009T | 0 | 0 | 0 |
| MEW003 | 0 | 0 | 0 |
| SH001 | 0 | 0 | 0 |
| MEW002 | 0 | 0 | - |
| 51148 | 0 | 0 | 0 |

We investigated the ancestry structure of the NYC coyotes and found that several NYC coyotes carry dog ancestry blocks on Chr 6 (Fig. 4). Two coyotes lacked dog ancestry on their chromosome 6 (MEW003 and SH002). We annotated 55 ancestry blocks of either homozygous dog or heterozygous coyote-dog ancestry (Table 4). Of the coyotes with dog ancestry, NY05 carried 14.9 times more dog ancestry in homozygous blocks than in the heterozygous configuration (41 and 2.7 Mb, respectively). Coyote NY04 similarly carried 41.8 Mb of dog ancestry in homozygous blocks but with 10.2 Mb of dog ancestry found in heterozygous blocks. Male coyote NY01 also carried a large fraction of dog ancestry on chromosome 6 in homozygous blocks (32Mb) paired with 24Mb of dog ancestry situated in heterozygous blocks. We also found 11 dog ancestry blocks in the region of chromosome 6 that contains the human-directed hypersociability TE alleles (Table 4). For the three coyotes that have the TE alleles, MEW001 and NY04 carried 100% dog ancestry while coyote NY01 carried 72.1% dog ancestry, which is independent evidence that these TE alleles were inherited from past interbreeding with dogs.

**Table 4.**
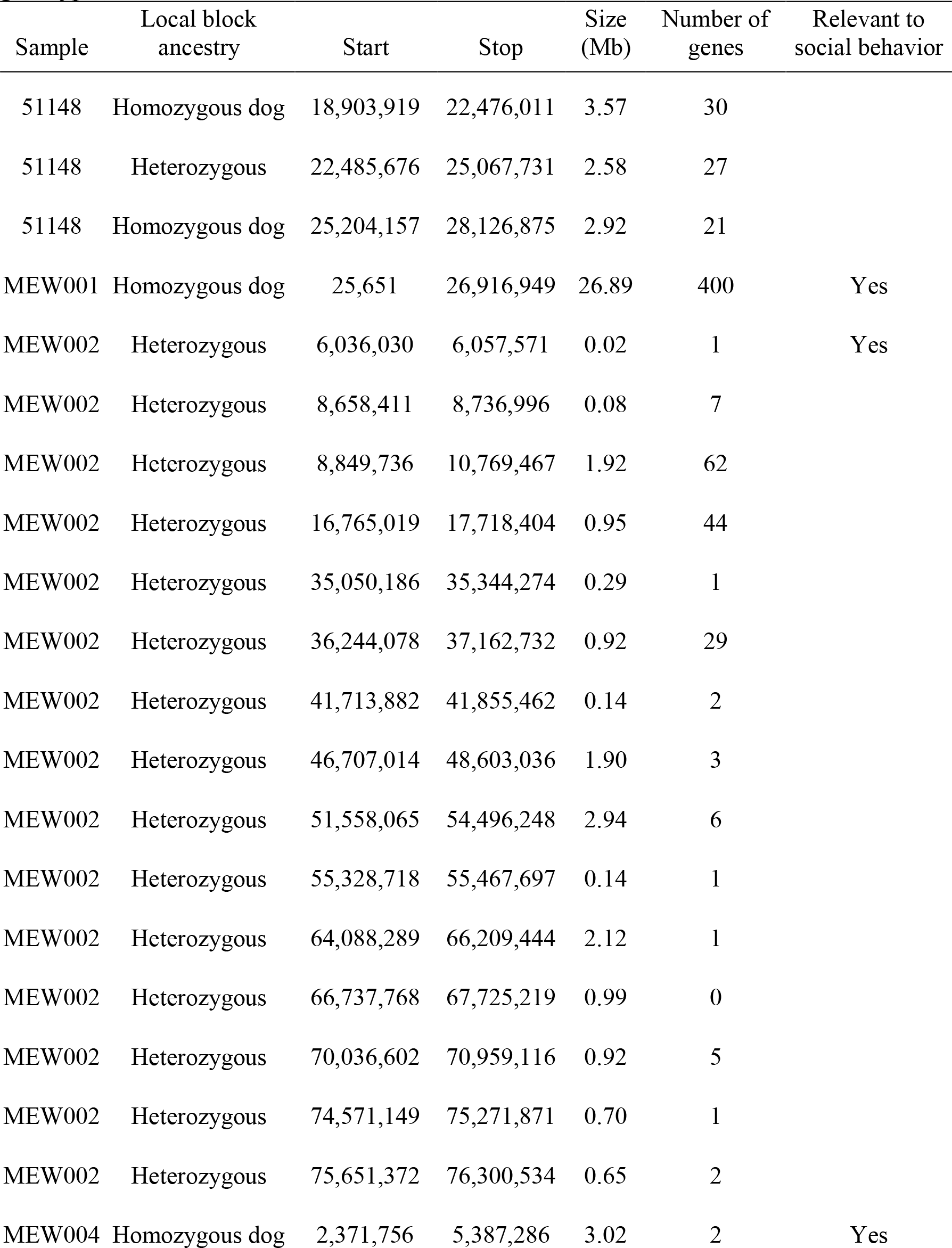
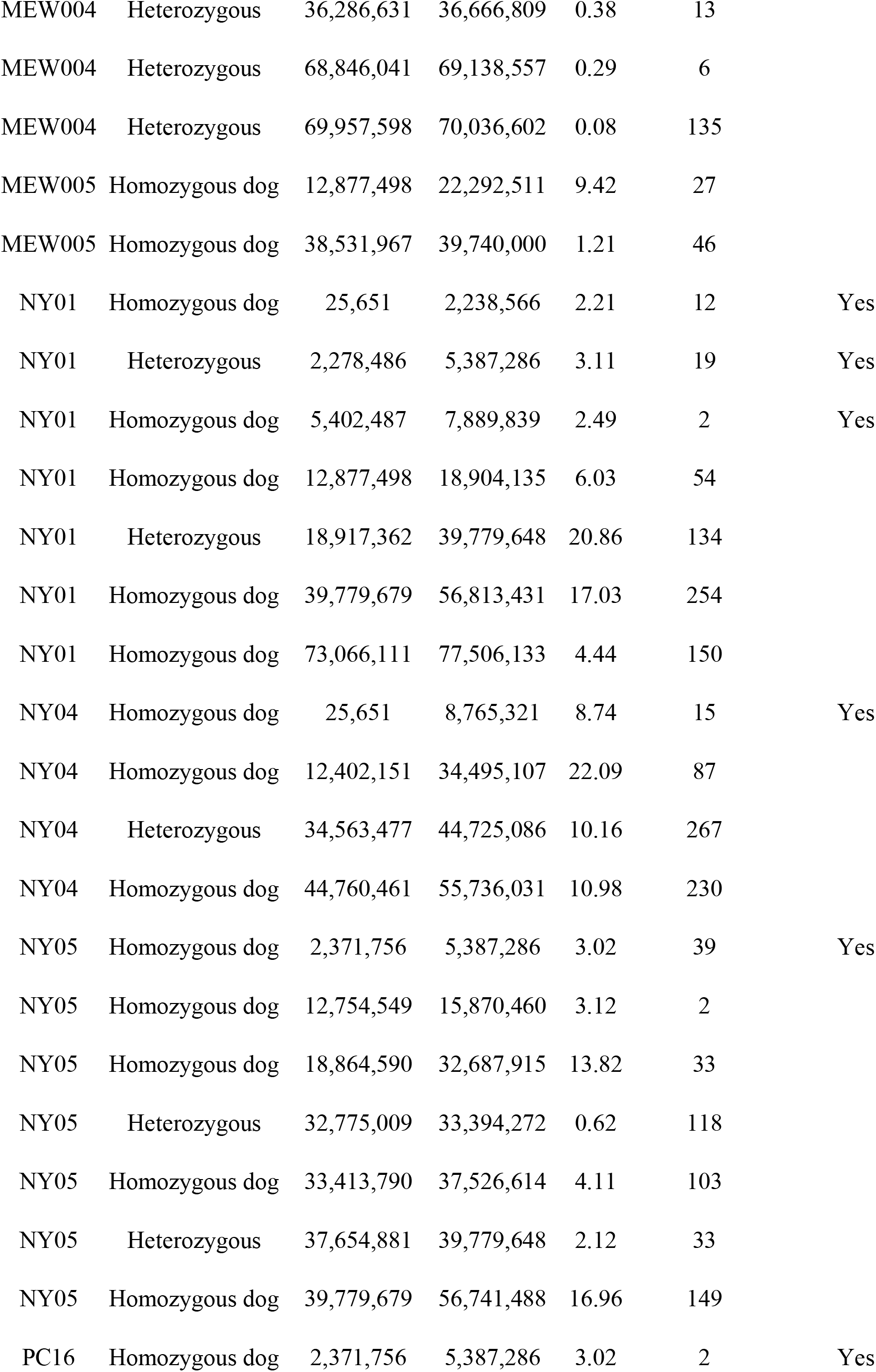
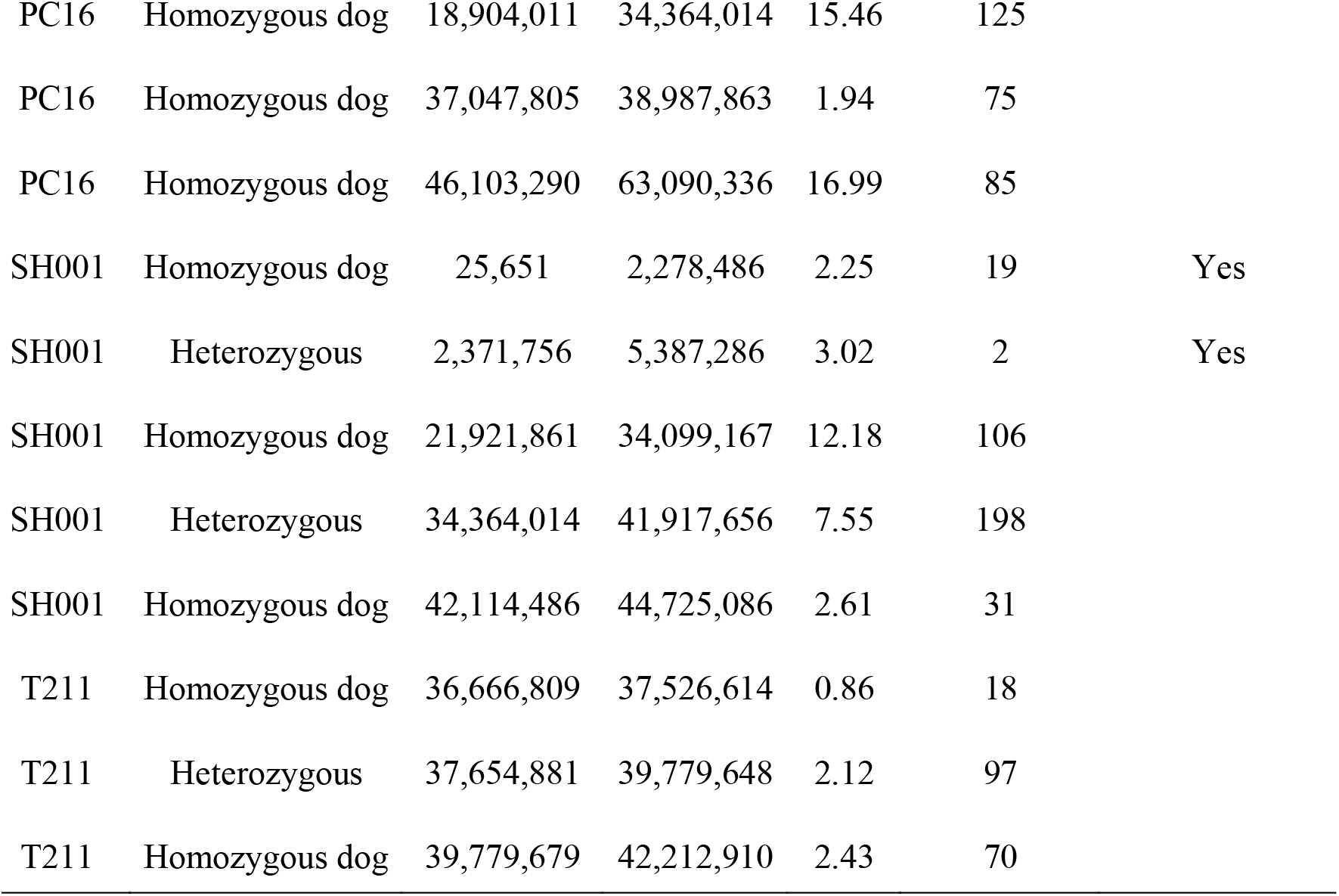
Genomic coordinates in canfam3.1 reference genome for blocks identified in the NYC coyotes with either homozygous dog or heterozygous coyote-dog ancestry. The number of genes found within each ancestry block is indicated, as well as if that block contains the TE alleles genotyped in Table 3.

**Figure 4.**
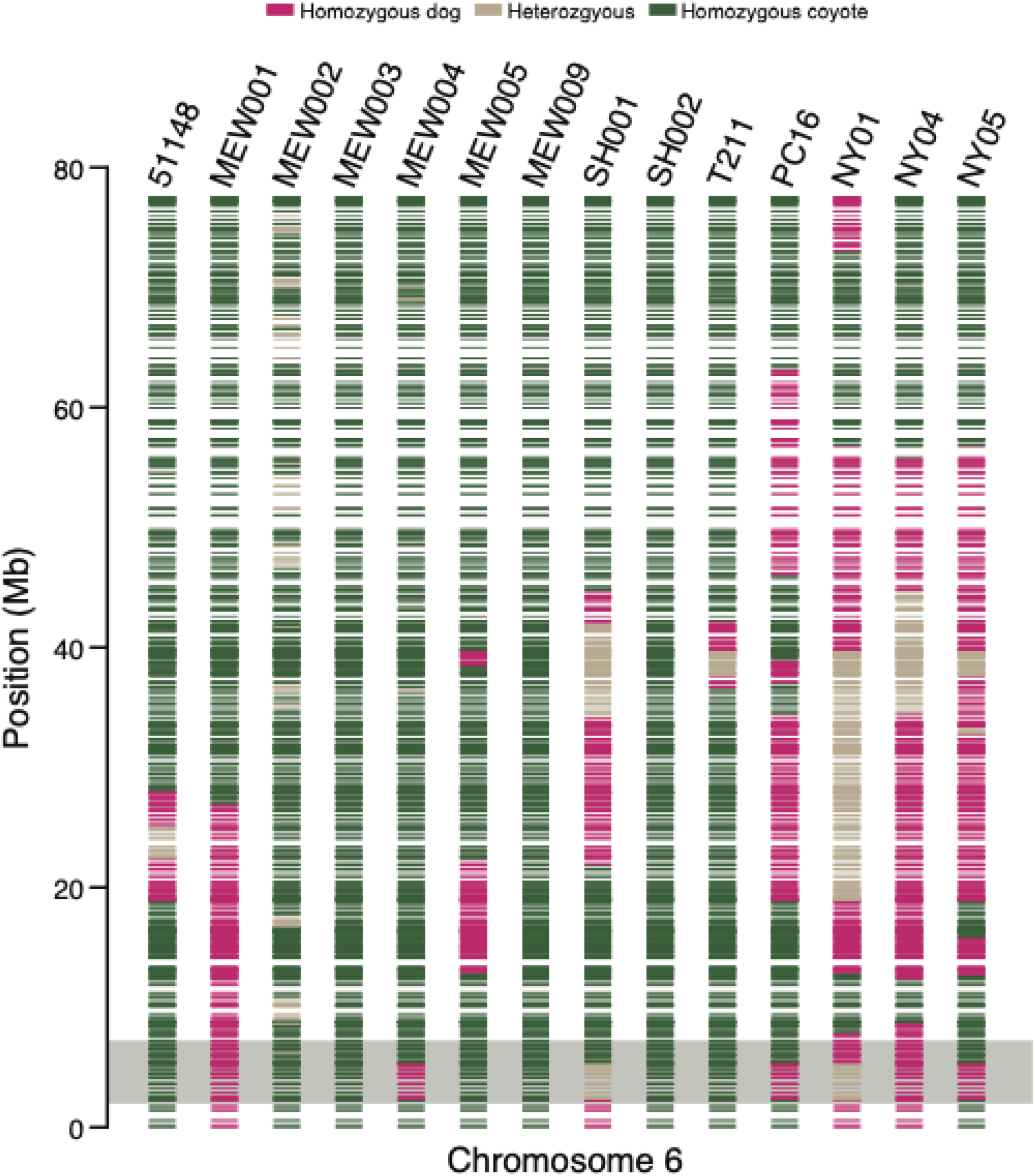
Ancestry blocks for canine chromosome 6. Each horizontal band is an ancestry block with the color of the block indicative of the ancestry state (homozygous or heterozygous) for the respective species identity. The chromosomal region highlighted by the gray box indicates the location of the hypersociability transposable element alleles.

## Discussion (word count: 1087)

We analyzed genome-wide SNP genotype data and found that NYC-area coyotes carry, on average, low levels (5-10%) of domestic dog genetic ancestry. Further, we provide the first documented evidence of first- and second-generation coyote-dog hybrids in NYC. The F1 coyote-dog hybrid is an adult male (NY01) of the Elmjack group and was mated with an unrelated female mate (T211) who carried only background levels of dog introgression. We further documented close genetic relationships (pairwise r = 0.35–0.5) of these adult animals with two Elmjack pups (NY04 and NY05). Each pup notably carried 25% dog genetic ancestry, the expected proportion for offspring of a cross between a coyote-dog F1 hybrid (NY01) and a non-admixed coyote (T211). Together with direct observations, we conclude that NY01 and T211 were a breeding pair and NY04 and NY05 were their offspring.

We found several animals in our NYC regional sample set that were more distantly related (e.g., second degree relatives). Our overall findings support Henger et al. (2020) who concluded that this population descends from a small number of founders and remains relatively small given the kinship found among multiple individuals. We also found evidence that coyotes disperse across habitat patches in the metropolitan landscape, at distances comparable to the species’ known dispersal abilities (Harrison 1992, Way 2007), as we documented close kinship ties between animals across the region. The urban and island nature of NYC has somewhat delayed colonization of Queens and the rest of Long Island, but the Elmjack family and other reports (Nagy et al. 2017, Henger et al. 2020) show that dispersal is proceeding, nonetheless.

Studies across multiple species show that animals with frequent human encounters often exhibit reduced fear of, and habituation to, humans over time (Carrette & Tella 2013; Cook et al 2017; Martin & Realé 2008; Uchida et al 2017; Vincze et al 2016). For coyotes, there appears to be competing ecological costs and benefits to using anthropogenic resources. Being tolerant of human activity may allow movement across the “concrete jungle” and/or persistence in an area with high human activity. However, overly bold coyotes will ultimately cause conflict and be removed. Gehrt et al. (2011) posited that the coyote is a “misanthropic synanthrope”, arguing that successful coyotes are those that can take advantage of the urban landscape, but avoid interacting with humans directly.

Coyotes thus seem to be pulled in two directions regarding human activity, and how an individual coyote behaves depends greatly on what they experience. Schell et al. (2018) observed captive coyote pairs over successive litters and found that parents engaged in riskier behavior (i.e., foraged more frequently) with their second versus first litters, and that parental habituation may result in reduced fear of humans in their offspring. However, in field settings, urban coyotes usually avoid people spatially (Atwood et al. 2004, Gerht et al. 2009, Thompson et al. 2021) and temporally (Gese et al. 2012). Young et al. (2019) found that coyotes that were hand-fed were more likely to subsequently approach humans and were harder to re-condition towards avoidance, showing that loss of fear of humans and associated habituations are behaviors learned by individual coyotes. In other words, coyotes’ behavior regarding humans depends on humans’ behavior regarding coyotes.

A potentially key component of this interplay may occur at a genetic level. There is evidence that domesticated and derived traits can be transferred back into wild relatives, and possibly confer an adaptive advantage. For instance, the melanistic *K* locus mutation causes melanism in coat color in North American wolves and derives from past hybridization with domestic dogs. This mutation is found at a high frequency in forested habitats and exhibits a molecular signature of positive selection, indicating it confers an adaptive advantage (Anderson et al 2009). Similarly, reduced anxiety and fear towards humans may confer an adaptive advantage, at least to a point, to coyote individuals inhabiting and traversing human dominated landscapes.

Here, we examined three candidate alleles that are known to influence human-directed social behavior in canines (vonHoldt et al. 2017). These alleles are retrotransposons and their copy number increases sociability with humans. We wanted to determine if coyotes with higher dog genetic ancestry also potentially carried these derived alleles, which may explain the anecdotal information about the Elmjack coyotes as being more conditioned to humans. The dog-derived alleles segregate at high frequencies in dogs but is exceedingly rare, if not fully absent, in coyote populations (vonHoldt et al. 2017). In agreement with past surveys, we found that the majority of coyotes in our study carried no derived alleles at these loci. However, three coyotes were heterozygous for two of the three loci. Two of these coyotes were the putative father-offspring pair NY01-NY04 from Elmjack, which both have significant genome-wide dog ancestry, and exhibited unusual dog-like pelage patterns (Fig. 1).

Observing two heterozygous genotypes per coyote significantly increases the probability that they would show social behavior towards humans (vonHoldt et al. 2017). In fact, the Elmjack coyotes were euthanized due to human-coyote interactions, including observations of feeding by humans (Duncan et al. 2020). Diet analyses of the Elmjack coyotes found a high percentage of urban commensals (e.g. rats) and anthropogenic remains (e.g. foil, paper, plastic; Duncan et al. 2020). Given the preponderance of dog genetic ancestry concomitant with the diet analysis, these coyotes faced a difficult existence as their genetics may have predisposed them to human interactions in an environment where such interactions do not typically have a positive outcome for wildlife (Gompper 2002; Breck et al 2019; Young et al 2019). Interestingly, the third coyote to carry the derived alleles (MEW001) did not show any appreciable dog genetic ancestry. Although this pattern is seen in just one animal, it suggests that historical introgression from dogs into these urban coyotes may have an impact on behavioral traits.

Our genetic evidence of recent coyote-dog hybrids, the small population size, and the relatively high frequency of hypersocial alleles suggest that genetic monitoring will be critical to understand how to coexist with coyotes in the NYC area. Rapid genetic changes through occasional coyote-dog hybridization could lead to changes in genetic background and behavioral traits associated with urbanization (Henger et al 2019). The rate of F1 hybridizations may increase as coyotes enter eastern Long Island and have limited options for mates. Researchers should combine ecological and genetic studies to fully explore the trophic and human dimensions consequences of this ongoing range expansion.

## Data Availability

The 17 RADseq BAM files sequenced in this study have been submitted to the NCBI BioProject database (https://www.ncbi.nlm.nih.gov/bioproject/) under accession number PRJNA857904. See Table S1 for references to additional public data included in this study.

---

**Supplemental Figure S1.**
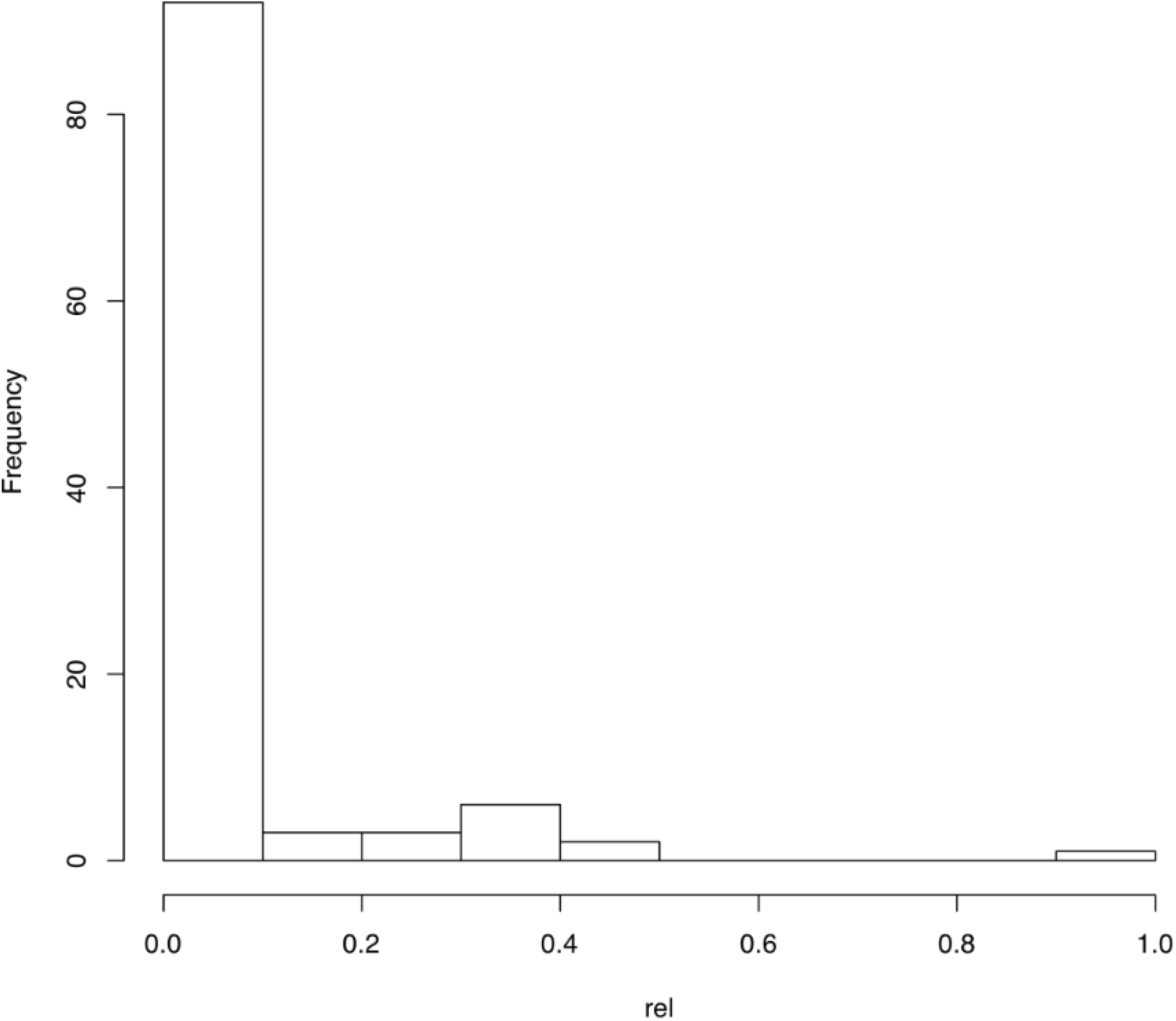
Histogram of pairwise relatedness values from the dyadic likelihood estimator of 15 target coyotes sampled in the NYC metropolitan area genotyped for 240 SNP loci genotyped.

